# Phase-dependent closed-loop modulation of neural oscillations in vivo

**DOI:** 10.1101/2020.05.21.102335

**Authors:** Colin G. McNamara, Max Rothwell, Andrew Sharott

## Abstract

Normal brain function is associated with an assortment of oscillations of various frequencies, each reflecting the timing of separate computational processes and levels of synchronization within and between brain areas. Stimulation accurately delivered on a specified phase of a given oscillation provides the opportunity to target individual aspects of brain function. To achieve this, we have developed a highly responsive system to produce a continuous online phase-estimate. In addition to stable oscillations, the system accurately tracks the early cycles of short, transient oscillations and can operate across the frequency range of most established neuronal oscillations (4 to 250 Hz). Here we demonstrate bidirectional modulation of the pathologically elevated parkinsonian beta-band oscillation (around 35 Hz) in 6-OHDA hemi-lesioned rats. Beta phase, monitored using a single channel electrocorticogram above secondary motor cortex, was used to drive electrical stimulation of the globus pallidus on one of eight phases spanning the oscillation cycle. Stimulation of the early ascending phase suppressed the oscillation whereas stimulation of the early descending phase was amplifying. By implementing a rule that prevented stimulation when the phase estimate was unstable, we achieved a system that could adapt stimulation rate and pattern to respond to the changes produced in the target oscillation. This allowed the electronic system to create and maintain a state of equilibrium with the biological system resulting in continuous stable modulation of the target oscillation over time. These results demonstrate the feasibility of phase locked stimulation as a more refined strategy for remediation of pathological beta oscillations in the treatment of the motor symptoms of Parkinson’s disease. Furthermore, they establish the utility of our algorithm and allow for the potential to assess the contribution of rhythmic activity in neuronal computation across a number of brain systems.

## Introduction

Rhythmic activity as reflected in neural oscillations plays a fundamental role in normal brain processing coordinating activity within and across regions (Buzsaki and Draguhn, 2004, Fries, 2015). Such activities are often detected by recording a local field potential (LFP) or electrocorticogram (ECoG), which predominantly reflect synchronised synaptic activity between neurons in the vicinity of the recording electrode (Buzsaki *et al.*, 2012). As a result, the spike timing of specific cell types is organised around the phase of LFP oscillations of different frequencies, which therefore serve as a temporal reference frame for local activity (Klausberger and Somogyi, 2008). Long range connections lead to synchronisation of oscillatory activities across brain regions, extending this temporal framework to networks (Siegel *et al.*, 2012). Both local and network level oscillatory activities have been found to correlate or contribute to motor, sensory and cognitive processing (Engel *et al.*, 2001, Buzsaki and Draguhn, 2004, Fries, 2015). In line with these findings, sustained disruption or exaggeration of neuronal oscillations and the underlying synchrony contribute to many brain disorders (Uhlhaas and Singer, 2006, Brown, 2007).

To fully elucidate the role of neuronal oscillations in health and disease, it will be necessary to have tools that allow the selective manipulation of the strength and synchronisation of these activities. Many investigators have attempted to examine the effects of disrupting oscillations in a single brain area using pharmacology or electrical/optical stimulation. However, as most oscillatory activities extend across different brain areas, disruption of a single engaged node can permanently suppress oscillatory activity across the entire network. Alternatively, a common approach has been to try to increase oscillatory activity by stimulating continuously at a particular frequency. However, externally paced stimulation of a node – even if it successfully elicits responses in neurons of the target area – may not engage the network in the same way as stimulation timed to the physiological rhythmicity intrinsic to the network (Pogosyan *et al.*, 2009, Syed *et al.*, 2012). For example, externally paced stimulation can result in transient epochs of suppression or amplification depending on the relationship between stimulus train and the intrinsic rhythm (Cagnan et al., 2013, Holt et al., 2019). Moreover, either widespread disruption or continuous stimulation at the frequency of interest can have less specific effects on neuronal activity, affecting measures such as firing rate. An ideal perturbation would allow a given activity to be rapidly amplified or suppressed near instantly when required for the specific targeted intervention and be reliably held in that state for as long as is required.

An optimal solution to these challenges is to deliberately harness the intrinsic temporal dynamics of a given activity by applying synchronized stimulation to a specific *phase* of the ongoing LFP or ECoG oscillation (Rosin *et al.*, 2011, Holt *et al.*, 2016, Cagnan *et al.*, 2017, Widge and Miller, 2019), producing extended periods of amplification or suppression by consistently further energising or dampening the oscillatory activity. Such *phase-dependent* perturbation may also be effective while delivering lower total stimulation powers leaving non-oscillatory parameters of neuronal activity (e.g. firing rate) relatively undisturbed, at least when calculated over timescales that are longer than the oscillation cycle. Maximising the effects of such phase-dependent stimulation necessitates highly accurate closed-loop stimulation, whereby the instantaneous phase of the target oscillation is tracked online and used to drive the stimulus train. Several studies have provided proof-of-principle evidence that such an approach can be used to modulate the amplitude of ongoing neuronal oscillations (Nicholson *et al.*, 2018, Kanta *et al.*, 2019, Peles *et al.*, 2020).

One of the clearest translational applications for a phase-dependent approach is the disruption of beta oscillations in Parkinson’s Disease (PD). Following significant loss of dopaminergic innervation of the striatum, the spiking of neurons across the cortex, thalamus and basal ganglia oscillate at beta frequency (12-35 Hz) (Brown *et al.*, 2001). This rhythmic spiking is highly synchronised within and between brain areas, and is reflected by increases in the power and coherence of LFP oscillations across the circuit (Brown *et al.*, 2001, Williams *et al.*, 2002, Sharott *et al.*, 2018). The magnitude and/or synchrony of these network oscillations is correlated with the severity of motor symptoms and their improvement following treatment (Kühn *et al.*, 2006, Sharott *et al.*, 2014, Neumann *et al.*, 2016). In PD patients, continuous stimulation at a frequency that matched the patient’s individual peak beta frequency but independently drifting in phase alignment, has been used to demonstrate that three or more consecutive electrical pulses falling on a specific phase of the subthalamic nucleus LFP can temporarily suppress the magnitude of oscillations both locally and in cortex (Holt *et al.*, 2019). Using a closed-loop approach to consistently deliver stimulation on a given phase of the network oscillation has the potential to modulate ongoing beta oscillations over longer periods of time, and consequently reduce akinetic/rigid symptoms.

While proof of principle exists, determining the real potential of phase-locked stimulation has been limited by the technical challenges in realising accurate continuous phase targeting. This is particularly true in the case of many physiological oscillations that are constantly fluctuating in amplitude over short time scales, as is the case with Parkinsonian beta oscillations. In fact, transient episodes of elevated beta-band power are often referred to as “beta bursts” (Feingold *et al.*, 2015, Tinkhauser *et al.*, 2017). Here we overcame these difficulties by developing and deploying a novel, computationally lightweight and highly responsive approach to phase tracking in real-time. We implemented the approach as a hardware description in Verilog and it can be deployed in any recording system where electrophysiological data is routed through a field programmable gate array (FPGA), making it widely applicable. Using this approach, the phase of an ongoing beta oscillation, monitored using a simple electrocorticogram (ECoG), was used to accurately drive basal ganglia stimulation separately to eight evenly spaced phases spanning the oscillation cycle. Overall, the results show that phase-dependent stimulation can produce powerful, bidirectional modulation of constantly fluctuating ongoing network oscillations. Furthermore, we provide evidence that this modulation of amplitude resulted from entering a stable closed-loop state between brain and system.

## Results

We developed a novel approach to provide a continuous phase estimate in real time with near zero lag capable of delivering phase-dependent perturbations of neuronal activity. The approach has the potential to be applied widely in research and clinical domains. The computational resources required are minimal allowing it to be implemented as a digital circuit and we deployed this circuit to the field programmable gate array (FPGA) on the Intan electrophysiological recording system (http://intantech.com/). In contrast to some previous approaches (Cagnan *et al.*, 2017, Nicholson *et al.*, 2018, Zanos *et al.*, 2018), the transformation was performed in such a way that the estimate was produced instantly (within microseconds) of the corresponding signal sample being available. This online estimate was used to trigger stimulation at one specific phase (the target phase).

To demonstrate the efficacy of our system and to investigate its ability to modulate oscillatory activity, we performed phase-locked stimulation of the pathologically-exaggerated beta oscillations that occur in the parkinsonian brain (Hammond *et al.*, 2007). To this end, we performed recording and stimulation experiments in 6-Hydroxydopamine (6-OHDA) hemi-lesioned rats, which recapitulate these activities across the cortico-basal ganglia network. As in previous studies activities (Delaville *et al.*, 2015, Brazhnik *et al.*, 2016), these animals displayed a prominent spectral peak in the ECoG at the high beta range (~35 Hz), recorded while the animals were free to explore an open field (Fig. 1A). The phase of the ECoG beta oscillation was tracked and used to control the timing of electrical stimulation (50-70 μA) delivered to the globus pallidus (GP). We chose the GP as the stimulation target because of its key role in the generation of pathological beta oscillations in this model (Mallet *et al.*, 2008, Cagnan *et al.*, 2019, Crompe *et al.*, 2020). By measuring the impact of GP stimulation on the spectral properties of the cortical input signal, we were able to investigate the effects of phase-dependent perturbations of a synchronised network. All experiments were carried out with animals freely moving in an open field

**Figure 1.**
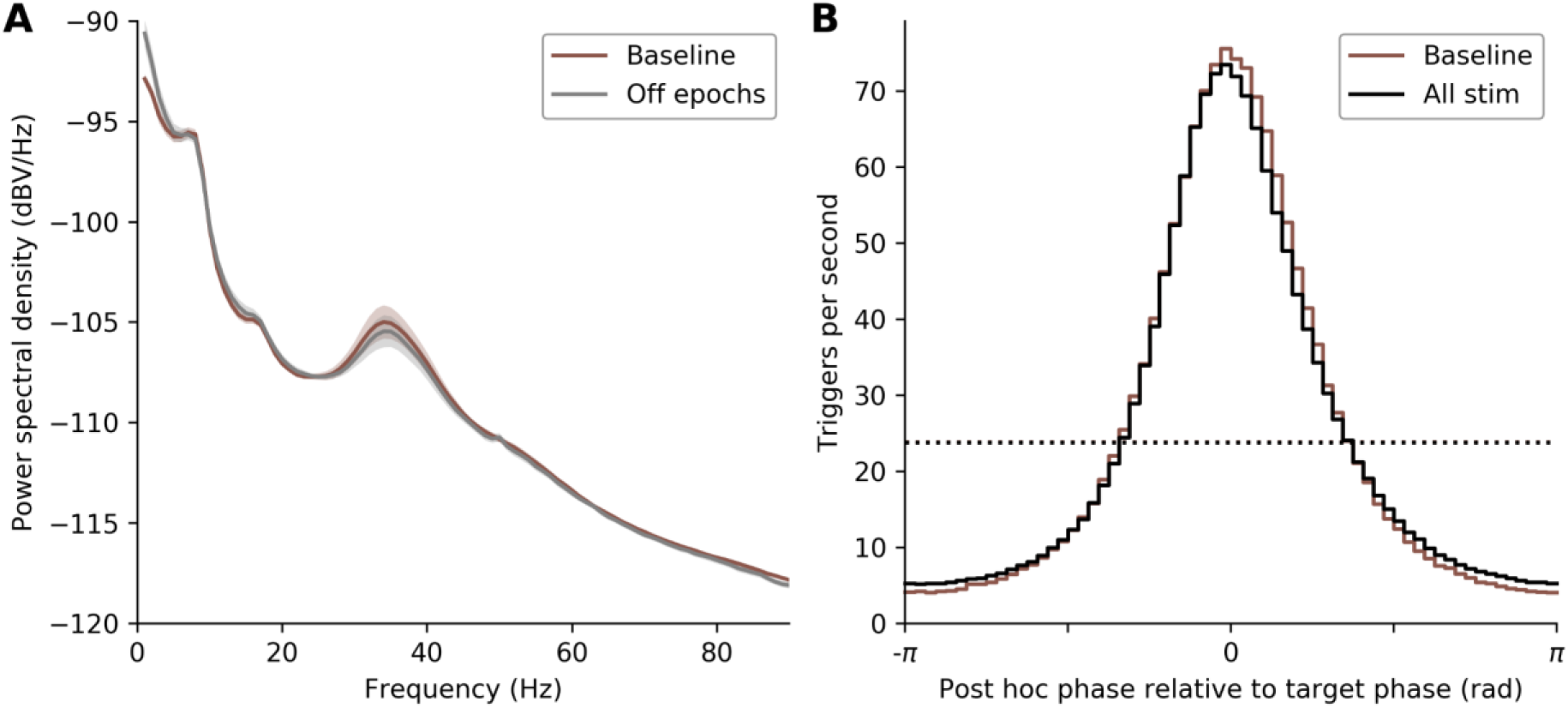
Phase-dependent stimulation targeting parkinsonian beta oscillations. **(A)** Power spectral density plot (mean ± SEM, n = 7 rats) showing elevated beta-band power in the absence of stimulation recorded from frontal cortical ECoG screws located ipsilateral to the 6-OHDA lesion. Elevated beta-band power was present in short (5 s) epochs with stimulation intentionally disabled embedded in stimulation enabled recordings (gray). The power in these epochs was very similar to that in longer baseline recordings without stimulation (brown). **(B)** Trigger accuracy histogram of ECoG phase calculated post hoc plotted relative to target phase. The accuracy and mean trigger rate (dotted lines) of the real time generated triggers, without stimulation enabled (brown) and with stimulation delivered to the globus pallidus (black) were almost identical, demonstrating that the system was able to maintain full accuracy in the presence of electrical artefacts and physiological perturbation produced by stimulating.

### High precision real-time phase tracking with and without delivery of electrical stimulation

We first evaluated the ability of our closed-loop system to accurately target 8 phases distributed evenly across the cortical beta oscillation cycle with and without the delivery of electrical stimulation to the GP. For each target phase, we made two types of recording. Firstly, in each animal we recorded with real-time phase tracking enabled and stimulation disabled (baseline recordings). This allowed us to define the spectral properties of the ECoG signal during spontaneous activity and record the real time generated triggers, when stimulation would have been delivered, without influence of physiological perturbation and/or stimulation artefact. Secondly, we performed recordings with closed-loop stimulation enabled (stimulation recordings). To assess the impact of stimulation on the ECoG, stimulation triggers were limited to 20 second epochs (on-epochs) separated by 5 second trigger free epochs (off-epochs), enabling us to compare epochs that were likely recorded in the same behavioural/arousal state. The beta-band spectral peak during stimulation free off-epochs was similar to baseline recordings (Fig. 1A), demonstrating that off-epochs were representative of general spontaneous activity.

We evaluated the accuracy of phase-dependent stimulation by computing histograms of post-hoc phase (calculated using a band-pass filter and Hilbert Transform) relative to intended real-time target phase (Fig 1B). To determine the overall accuracy of the system to trigger stimulation at any specific phase, we pooled the data across all target phases (the same analyses for specific phases are described below). Using this pooled data, 60.78 ± 4.28 % (mean ± SD, n = 7 rats) of the stimuli were delivered within a quarter of a cycle of the target phase. When the same analysis was performed using the baseline recordings (i.e. real-time phase tracking with stimulation disabled), the distribution of trigger phases with respect to the target phase was almost identical to that when stimulation was applied (Fig 1B, p = 0.14, Kolmogorov-Smirnov test, KS = 0.20). Thus, without considering the specific target phase, the accuracy of phase tracking was unaffected by stimulation.

In addition, the rate of real-time triggers in baseline recordings (23.92 ± 2.37 Hz) and stimulation recordings (23.78 ± 2.66 Hz) were not different (p = 0.55, paired t-test, t_6_ = 0.64) and were considerably less (31 %) than the theoretical maximum trigger rate (the centre frequency selected for phase tracking). The disparity was not due to the presence of off-epochs, as the trigger rate calculations were limited to on-epochs. This demonstrated a specific aspect of our approach, whereby delivery of stimuli was reduced during periods of low beta-band power when the phase estimate became unstable (see Methods). Overall, the real-time phase tracking approach employed here was able to track the ongoing phase of beta oscillations with the same accuracy during stimulation as when it was absent, and the system delivered the majority of stimuli in close proximity to the target phase.

### Phase-dependent modulation of beta power

We next addressed whether consistent stimulation at specific phases could modulate the power of the ongoing oscillation. As previously reported in this model (Cagnan *et al.*, 2019), the amplitude of beta in the raw ECoG recordings waxed and waned, leading to short periods of high amplitude (200-400 ms), often referred to as “beta bursts” (Tinkhauser *et al.*, 2017, Fig. 2A). For a given animal, some target phases appeared to reduce beta-band activity across the recording (Fig. 2B) while stimulation at other phases achieved the opposite. (Fig. 2C). These effects were particularly clear when the signals were filtered around the centre frequency used for phase-tracking (Fig 2 lower traces of each panel). Notably, the presence of regular pulses of stimulation at beta frequency did not always manifest as an increase in beta amplitude in the ECoG, suggesting that modulation of oscillation power was not due to stimulation artefacts.

**Figure 2.**
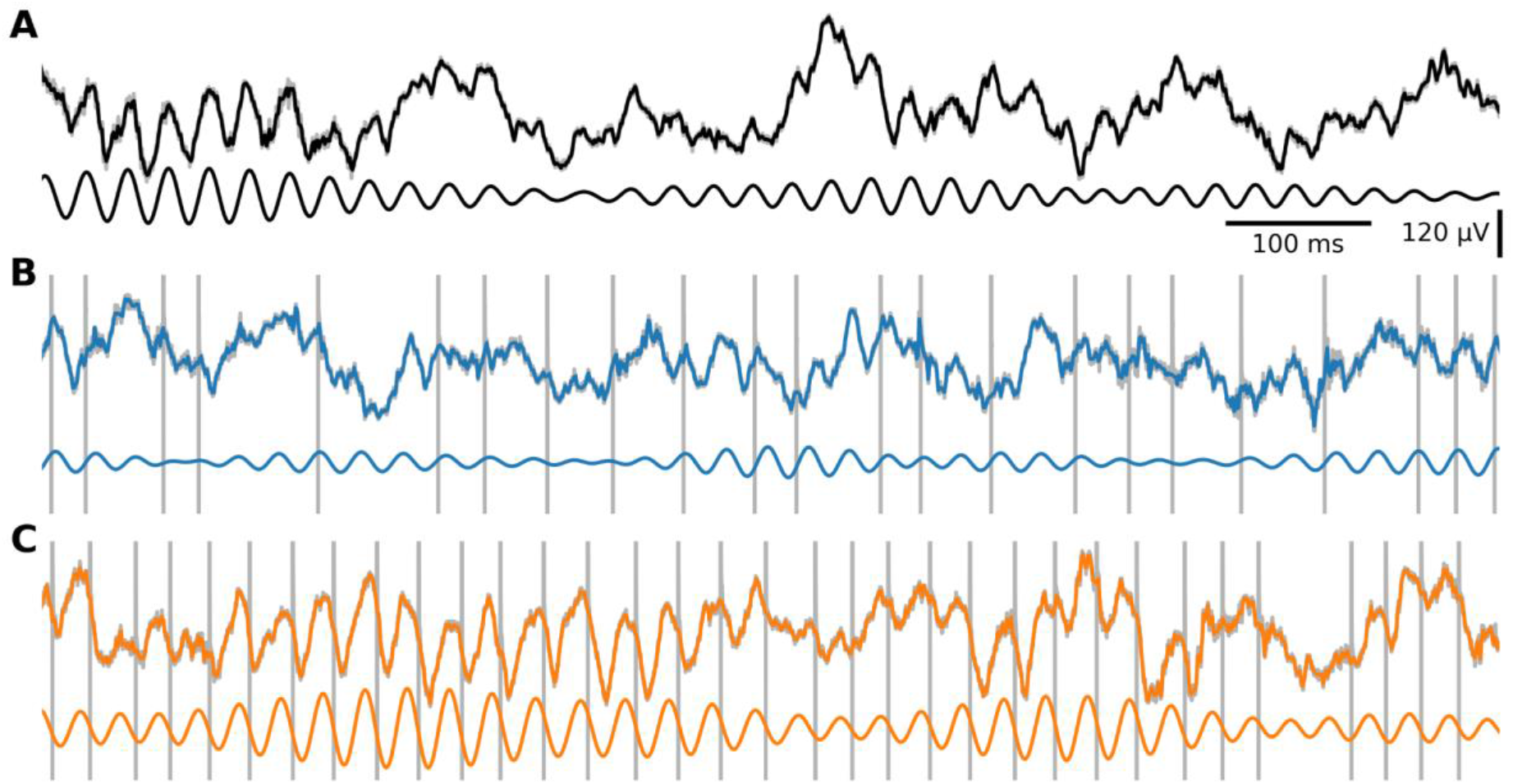
Example ECoG traces with amplification and suppression of beta-frequency oscillations. Example one second duration typical ECoG traces with **(A)** no stimulation, **(B)** stimulation delivered to the globus pallidus targeted to a suppressing phase of the ECoG and **(C)** stimulation targeted to an amplifying phase. Each panel shows the wideband signal downsampled to 1 kHz after removal of stimulation artefacts (coloured) superimposed on top of the original recording sampled at 20 kHz (grey). The stimulation artefacts visible as grey vertical lines are truncated in amplitude to aid visualisation of the physiological activity. The artefact free signal, filtered in the beta-band post hoc is shown below each trace. Note the mean beta-band amplitude compared to the no stim condition is reduced with suppressing stimulation and increased with amplifying stimulation.

To quantify these phenomena, we compared the difference in power spectral density (PSD) in the beta-band (centre frequency ± 5.5 Hz) during on-epochs and off-epochs for each stimulation recording at 8 target phases. The differences in beta power across these 8 phases were aligned relative to the target phase that led to the largest decrease beta power in that animal (suppressing target phase). The modulation of beta power was approximately sinusoidal, with a monotonically increasing progression of power to the target phase with the largest increase in spectral power (amplifying target phase) approximately half a cycle from the suppressive phase before monotonically decreasing back around to the suppressing phase (Fig. 3A). Absolute target phase significantly modulated beta-band power (p = 0.0013, one-way ANOVA, F_7, 48_ = 4.12) and the absolute phase of maximally suppressing and maximally amplifying modulation was different (p = 0.0022, Watson-Williams test, F_1, 12_ = 15.09). As depicted in Figure 3B, maximally suppressing modulation (−1.41 ± 1.08 dB) occurred around the early-ascending phase (1.36 ± 0.39 π rad) and maximally amplifying modulation (2.32 ± 0.80 dB) occurred around the early-descending phase (0.28 ± 0.32 π rad). To further visualise the modulation of beta-band power, we calculated the mean PSDs in on-epochs and off-epochs during stimulation recordings across animals for suppressing (Fig. 3C) and amplifying (Fig. 3D) target phases. Stimulation at the suppressing target phase flattened the beta-band spectral peak that was present in off-epochs, (Fig. 3C) whereas stimulation at the amplifying target phase led to large increase in size of the beta-band spectral peak (Fig. 3D). Taken together, these results demonstrate that both suppression and amplification of beta power can be achieved using GP stimulation timed to specific phases of the ECoG oscillation and additionally, that suppression and amplification occurred on similar absolute target phases across animals.

**Figure 3.**
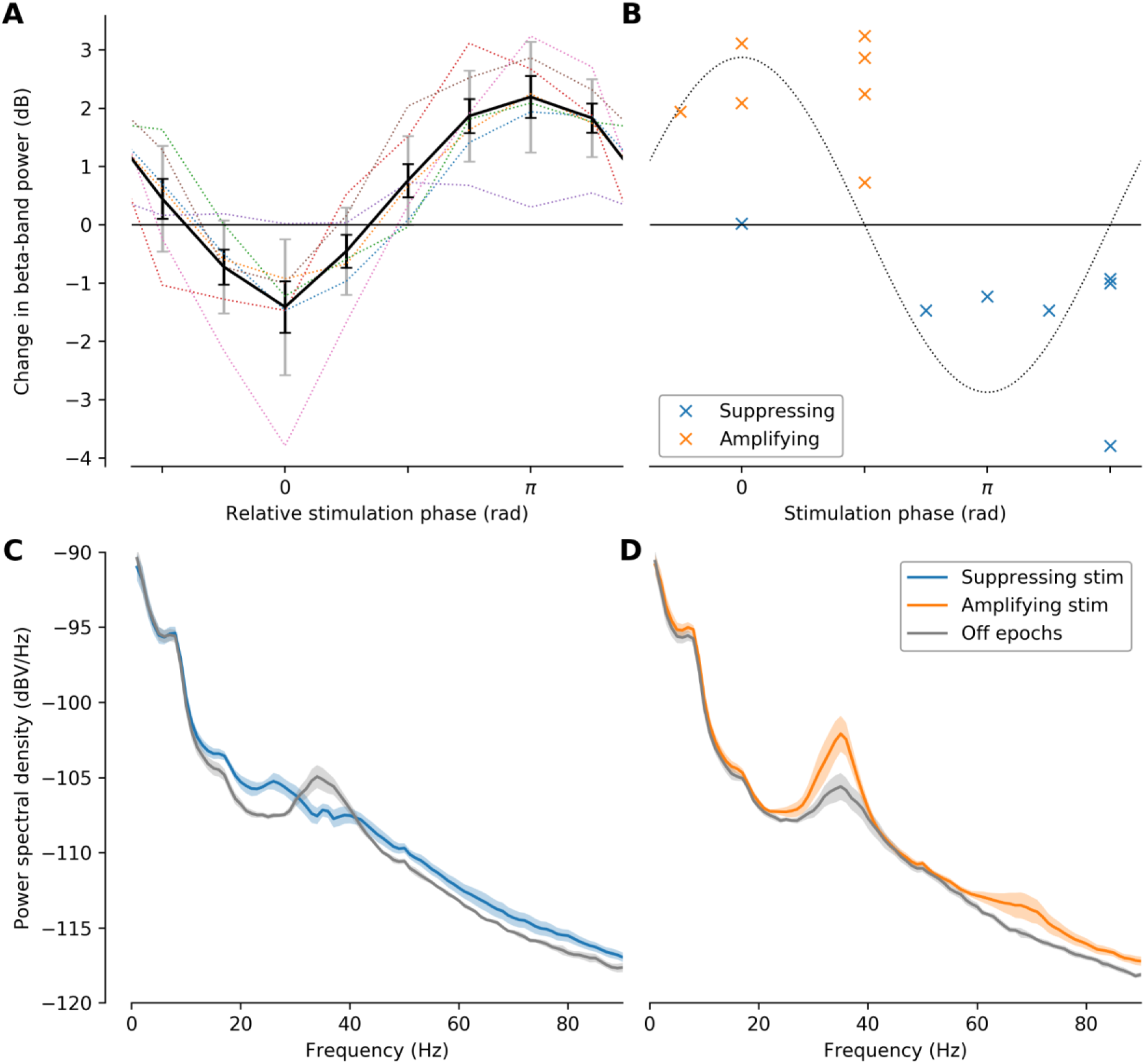
Phase-dependent modulation of parkinsonian beta-frequency oscillations. **(A)** Change in beta-band power due to stimulation across 7 rats for 8 target phases aligned by the most suppressing target phase for each rat. The change in beta-band power for each recording was calculated in dB as the ratio of the mean beta-band power in repeated short (5 s) epochs of no stimulation to the mean beta-band power in the longer intervening (20 s) epochs with stimulation enabled. The dotted lines show the values for each rat with the mean ± SEM in black and standard deviation in grey. **(B)** Absolute phase versus beta-band power of the most suppressing and most amplifying target phases for each rat. The dotted line provides a visual aid showing the location of the peak and trough of the beta-cycle. **(C, D)** Power spectral density plots (mean ± SEM, n = 7 rats) showing **(C)** a reduction in the beta-band spectral peak when stimulation was applied to the most suppressing target phase (blue) compared to stimulation disabled short epochs embedded in the same recordings (grey) and **(D)** an increase when stimulation was applied to the most amplifying target phase.

### Continuous phase-dependent perturbation leads to stable closed-loop states

Previous closed-loop methods of remediating pathological beta activity have focussed on delivering high frequency stimulation reactively when the amplitude of the oscillation breaks a pre-defined threshold (Little *et al.*, 2013). Stimulation results in attenuation of the length of these periods, bringing the amplitude below the threshold for stimulation faster than would occur at rest (Tinkhauser *et al.*, 2017), after which the process begins again. In contrast, our implementation of phase-dependent stimulation interacts continuously with the ongoing signal, dampening or energising the biological activity by stimulating once per cycle. With such a continuous approach, stimulation must adapt in response to changes in brain state and importantly it must respond to, and hence reflect, the change it seeks to create. Here, this was achieved by preventing stimulation when the phase estimate was deemed unstable (potential trigger events were supressed if it had been less than 0.8 of a target frequency period since the previous potential trigger event, see methods). This allowed the system to reduce the amount of stimulation when maintaining suppression of the target oscillation and increase the amount of stimulation when maintaining amplification of the target oscillation. The continuous modulation of beta amplitude is evident in the traces in Fig. 2 along with the differing properties of the stimulation patterns that arose by simply targeting different phases.

To assess the adaption of stimulation across the full data set, we analysed the phase-accuracy histograms and the mean stimulation rate separately for each condition (Fig. 4). Amplifying stimulation in contrast to suppressing stimulation had a higher mean stimulation rate (Fig. 4A, amplifying 25.75 ± 4.06, suppressing 21.19 ± 2.33, p = 0.0045, paired t-test, t_6_ = 4.42) and a greater accuracy measured as percentage of stimulation within a quarter cycle of the target phase (amplifying 74.96 ± 6.97 %, suppressing 47.29 ± 7.00 %, p = 0.0003, paired t-test, t_6_ = 7.26, see also stimulation accuracy histograms Fig. 4A). Since the absolute phase of amplifying and supressing stimulation was similar across animals (but not identical see Fig. 3B), these differences are also present in the mean trigger accuracy histograms across animals grouped by target phase (Fig 4B). Importantly, the differences in the stimulation produced by the electronic system – arising from the effect of the stimulation on the biological system (i.e. the modulation of beta-band power) – demonstrated that the electronic and biological systems entered into a combined closed-loop state, producing activity in each that would not have been possible in either, without the presence of the other.

**Figure 4.**
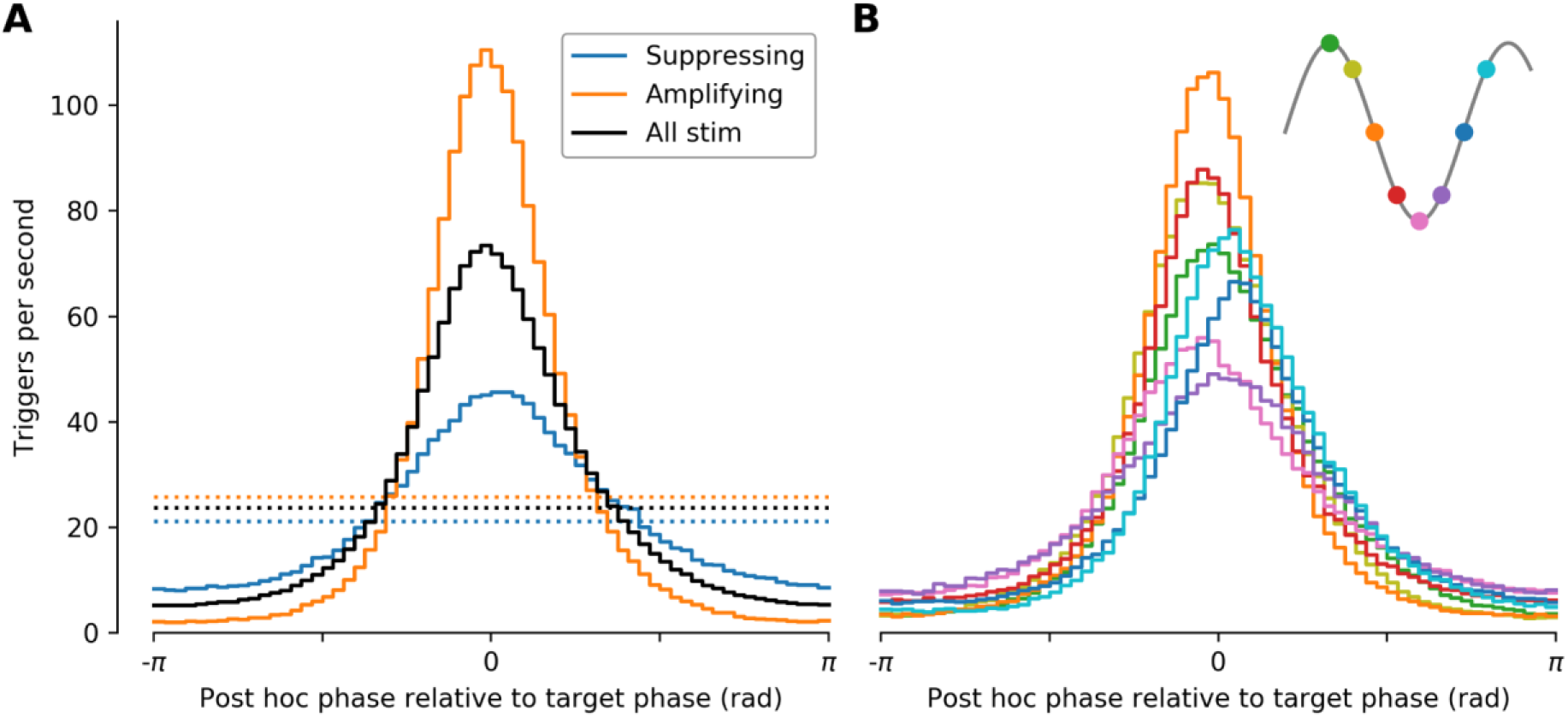
Stimulation properties reflected the direction of beta power modulation. **(A)** Trigger accuracy histogram of ECoG phase calculated post hoc plotted relative to target phase. Stimulation targeted to the most amplifying phase (orange) produced a higher rate of stimulation on the correct phase (0 rad) and a higher mean stimulation rate (dotted lines) compared to the mean across all target phases (black). Stimulation targeted to the most suppressing phase (blue) led to reduced accuracy and a lower mean stimulation rate (dotted lines) reflecting the systems success in suppressing the oscillation. **(B)** Mean trigger accuracy histograms for each target phase across all 7 rats plotted relative to target phase with inset showing the target phase of each trace.

## Discussion

Phase-dependent stimulation has widespread applications in fundamental and clinical neuroscience. Such applications will require adaptable systems that provide accurate phase tracking capable of functioning in the presence of stimulation. Here we demonstrate a self-contained system that, by meeting these criteria, can suppress or amplify the power of pathophysiological network oscillation. The direction of spectral modulation modified the continuous temporal dynamics of the stimulation itself, suggesting that the electronic system entered into a closed-loop steady state with the brain, rather than acting episodically in response to changes in signal strength before waiting for clear signal to re-emerge.

### Accurate Phase Targeting

A reasonable question to ask of any phase-locking application is how accurately the system is able to trigger stimulation at a specific phase of the target oscillation. However, accuracy of phase-tracking is not trivial to assess, as the outcome will depend on several factors including the signal-to-noise ratio of the input signal, the post-hoc method used to assess the phase at which the stimuli were delivered, the wider influence of stimulation and decisions around stimulus artefact removal. The post-hoc calculation of phase, in comparison to the online estimate, is based on the data before and after a given data point and is thus, by definition, a different measure to online phase and unlikely to exactly correspond. Furthermore, there is not a single correct phase estimate as measuring one of instantaneous phase, amplitude or frequency of an imperfect periodic process requires assumptions about the others. This forces choices which are context and application specific. We assessed the phase of stimulus delivery with a Hilbert-transform based method as this has been widely used by us and others to define the phase of beta oscillations in recorded data. Under the conditions described, the phase (calculated post-hoc) of 60% of stimulation triggers was within a quarter of a cycle of the target phase, both when stimulation was applied and when it was disabled across a group of animals. Assessing the post-hoc phase of real-time triggers with stimulation disabled was an important control, as spectral bleed through from the artefact removal hold circuit (see methods) could potentially confound the phase estimate by introducing an oscillatory process to which the system could lock. The similarity of results across stimulation-enabled and -disabled conditions provides evidence that stimulation itself was unlikely to have contributed to phase locking accuracy.

A second important question is *how* to use the ongoing phase estimate to time stimulation. We used a small range of stimulation amplitudes and chose to deliver a maximum of one stimulus on any single cycle. This dictated that the maximum stimulus rate was the centre frequency of phase locking. However, an accurate phase-estimate could be used to deliver any number of stimulation protocols across the phase/amplitude space (Weerasinghe *et al.*, 2019). A novel feature of our approach was to prevent stimulation when the phase estimate became unstable. This occurred at times of lower oscillatory activity in the neuronal elements underlying the input signal causing the associated spectral component to fall below the noise floor created by the multitude of co-occurring neuronal activities. We reasoned that stimulating at these times would be counterproductive, as it would not be timed to optimally interact with the neuronal activity of interest and had the potential to produce undesired outcomes.

### Phase-dependent Modulation of Beta Power

The central finding of this study was that stimulation of the globus pallidus at specific phases of the ongoing ECoG beta oscillation could suppress or amplify the power of that input signal. To address the mechanisms behind this modulation, it is necessary to consider the neuronal circuitry and spontaneous temporal dynamics underlying the parkinsonian beta oscillation. In the 6-OHDA lesioned rat, pathologically enhanced beta oscillations are synchronised across every part of the cortico-basal ganglia-thalamic loop (Mallet *et al.*, 2008, Devergnas *et al.*, 2014, Brazhnik *et al.*, 2016, Cagnan *et al.*, 2019). Both physiological and computational studies suggest that the GP plays a particularly important role in propagating/maintaining beta oscillations around this circuit (Nevado-Holgado *et al.*, 2014, Corbit *et al.*, 2016, Cagnan *et al.*, 2019, West *et al.*, 2020). There are two main neuron types in the GP. Prototypic neurons send axonal projections to the subthalamic nucleus (STN) and basal ganglia output nuclei, whereas arkypallidal neurons send widespread axonal projections exclusively to striatum (Mallet *et al.*, 2012, Abdi *et al.*, 2015). In parkinsonian animals, the spiking of neurons in both populations is locked to the cortical beta oscillations, with mean phase angles that are approximately 180 degrees apart. Thus stimulation of the GP at a specific phase of the ECoG beta cycle could lead to several outcomes for spike trains of individual neurons and the temporal relationships between them. For example, if stimulation arrives during the spiking of local synchronised neurons, it could reinforce both the individual oscillation of single neurons and synchronisation between neurons of the same population. In contrast, if stimulation arrives at another specific offset from the rhythmic firing, it might disrupt the oscillation and desynchronise the population. As the pattern of collaterals between the two neuron types has not yet been defined, it is currently difficult to predict how the stimulation of one population will affect the other. In MPTP-lesioned primates, stimulation of the GP triggered by rhythmically firing motor cortical neurons at a fixed-delay suppresses pathological oscillatory activity and ameliorates parkinsonian motor deficits (Rosin *et al.*, 2011). While the phase of cortical activity was not explicitly tracked in that study, and stimulation was delivered to the internal segment of the GP, it seems likely that the effect was produced through similar mechanisms to our own set-up. Further recordings of the activity of specific populations, together with computational models will be needed to understand how phase-dependent perturbations modulate both local and network oscillations in this and other systems.

### Closed-loop dynamics

Closed-loop manipulations of the brain are in an early stage of development. Studies thus far have focussed on how the stimulation is influencing a particular parameter of brain activity. To our knowledge, little attention has been paid as to how any change in brain activity will in turn influence the dynamics of the stimulation. Our system prevented stimulation being delivered when the phase estimate was unstable, which resulted in a trigger rate around 10 triggers/s below the theoretical upper bound dictated by the pre-set closed-loop centre frequency parameter (f_c_). This allowed us to examine how stimulation at different phases influenced the trigger rate, which could change in either direction. Notably, stimulation at the suppressive trigger phase reduced the trigger rate, presumably because the reduction in oscillatory signal led to a larger amount of time where the phase estimate was unstable causing stimulation to be supressed by the system. In contrast, by reinforcing the oscillation and making the phase estimate more reliable, stimulation at the amplifying target phase led to a higher trigger rate. This interaction between the modulated neuronal signal and features of the closed-loop system thus produced a state of equilibrium allowing both the biological system and the electronic system to operate with stable characteristics only possible due to the presence of the other. The ability of a closed-loop system to produce and maintain a stable state of equilibrium with the biological system under modulation is likely to be particularly important in clinical applications of phase dependent stimulation, where stability is required over days and weeks.

### Applications in clinical and fundamental science

Several investigators have highlighted the potential of phase-locked stimulation for treating neurological and psychiatric disorders (Rosin *et al.*, 2011, Brittain *et al.*, 2013, Azodi-Avval and Gharabaghi, 2015, Meidahl *et al.*, 2017, Widge and Miller, 2019). Parkinson’s Disease provides one of the best examples of such application, with both tremor and beta frequency oscillations clearly associated with symptoms. Indeed, stimulation of cortex or thalamus timed using the phase of a peripherally recorded tremor oscillation was one of the first examples of the power of the approach to suppress and amplify ongoing oscillations (Brittain *et al.*, 2013, Cagnan *et al.*, 2017). In comparison to tremor oscillations, beta activity presents a different challenge, given that it necessitates central recording of the signal and that the oscillations themselves are far less consistent. In addition to our study, a recent preprint (Sanabria *et al.*, 2020) demonstrates similar phase-dependent modulation of beta oscillations in MPTP-lesioned primates, using the beta-phase of LFP recordings from the STN to trigger stimulation in the GPi. Beta oscillations in the parkinsonian primate have a lower centre frequency (8-15Hz) than in awake rodent and beta bursts last for several seconds (Deffains *et al.*, 2018), whereas those in rodents tend to be subsecond in duration (Cagnan *et al.*, 2019). In PD patients, while there is a wide range of centre frequencies, the average is closer to the low beta seen in primates than the higher beta frequency in awake rats. In terms of temporal dynamics, however, the subsecond beta bursts described in rodents more closely resemble those described in patients (Cagnan *et al.*, 2019). Thus, between this study and that of Sanabria and colleagues there is evidence that phase dependent stimulation can effectively modulate beta oscillations across a wide range of spatial, spectral and temporal parameters that encompass those described in patients. Notably, two separate studies using healthy primates have also demonstrated that phase-dependent stimulation of cortical beta oscillations can modulate plasticity (Zanos *et al.*, 2018) and behavioural performance (Peles *et al.*, 2020), demonstrating the utility of such neuromodulation.

Together with our own proof-of-principle demonstration of phase-dependent suppression of beta oscillations (Holt *et al.*, 2019), there is now ample evidence that this approach could provide a more refined strategy for modulating beta oscillations in PD patients than current interventions. Implementing closed-loop strategies is becoming increasingly tractable given the emerging generation of medical devices that can both sense and stimulate (Cagnan *et al.*, 2019). Approaches to overcoming the challenges of real-time phase estimation include future signal prediction using an auto regressive model to improve responsiveness and overcome the algorithmic delays introduced by applying a real-time band pass Hilbert filter (Chen *et al.*, 2011, Chen *et al.*, 2013, Blackwood *et al.*, 2018). Our approach offers a number of attractive features making it ideal for incorporation into future devices. It is extremely responsive; having minimal filter delay allows it to produce estimates relating to the current sample as opposed to a number of samples previous. This is because it does not rely on a traditional band pass filter to restrict the estimate to the frequency band of interest. It requires minimal computational resources allowing it to be implemented as a digital circuit (as was the case here) or adapted to execute on a low power microcontroller and still leave additional computational resources free. This has the related advantages of requiring little power to operate, the ability to compute each estimate near instantaneously and, while not demonstrated here, the ability to operate at centre frequencies spanning the full range relevant in neuroscience (4 to 250 Hz). Furthermore, that the algorithm is both capable of incorporation into implantable devices and research electrophysiology platforms makes it well suited to translatable studies.

While implementation of phase-loop stimulation has clear translational utility, there are even wider implications and applications of this approach for fundamental science (Siegle and Wilson, 2014, Nicholson *et al.*, 2018, Zanos *et al.*, 2018, Kanta *et al.*, 2019, Peles *et al.*, 2020). The long running debate as to how or whether neuronal oscillations contribute to computation has been hampered by the lack of tools to target oscillatory processes over other parameters of neural activity. Here we demonstrate that it is possible to cause subtle modulations in the power of neuronal oscillations within their dynamic range using low amplitude stimuli. Such approaches could be further refined using optogenetic stimulation to target specific neuronal populations. Utilising phase and frequency of input signals from one area to control the timing of stimulation or inhibition to another part of the network over a range of timescales presents an unprecedented opportunity to define the role of neuronal oscillations across different systems.

## Methods

### Subjects and surgical procedures

Experiments involving animals were conducted in accordance with the UK Animals (Scientific Procedures) Act 1986 under personal and project licenses issued by the Home Office following ethical review. Male Lister Hooded rats (starting weight 350 – 450 g) were housed with free access to food and water in a dedicated housing room with a 12/12-h light/dark cycle and underwent two separate recovery surgical procedures performed under deep anaesthesia using isoflurane (4% induction, 2-0.5% maintenance) and oxygen (2 l/min). Local anaesthetic (Marcaine, 2 mg/kg, 2.5 mg/ml) and non-steroidal anti-inflammatories (Metacam, 1 mg/kg, 5 mg/ml) were administered subcutaneously at the beginning of all surgeries, while opioid analgesia (Vetergesic, 0.3 mg/ml, 0.03 mg/kg) was also provided for three consecutive post-operative days. The first procedure produced a unilateral lesion of dopaminergic neurons of the substantia nigra pars compacta by intracranial injection of the neurotoxin 6-hydroxydopamine (6-OHDA) at their cell bodies through a glass pipette using stereotaxic coordinates 4.9 mm posterior and 2.3 mm lateral from bregma at a depth of 7.8 mm from the brain surface. 6-OHDA was dissolved immediately before use in phosphate buffer solution containing 0.02% w/v ascorbate to a final concentration of 6 mg/ml. Between 0.1 and 0.15 μl of 6-OHDA solution was injected at a rate of 0.01 μl/min through the pipette, which was left in place a further 5 minutes before being withdrawn. Following full recovery and not less than 13 days later, animals underwent a second procedure to implant pairs of stainless steel stimulation electrodes (California Fine Wire, stainless steel, bifilar, heavy formvar insulation, 127 μm strand diameter) and M1.4 cerebellar reference and frontal electrocorticography (ECoG) screws. Electrical stimulation was delivered to the globus pallidus through electrodes implanted using stereotaxic coordinates 1 mm posterior and 3.1 mm lateral from bregma at a depth of 6.4 mm from the brain surface. ECoG was measured from the most frontal screw located above motor cortex at approximately 4.6 mm anterior and 1.6 mm lateral from bregma. Neurotoxin injections, electrical stimulation and ECoG recording were all performed on the right hemisphere. During data acquisition which took place over multiple days, rats freely explored a 90 by 50 cm open field surrounded on five sides with electrical shielding located in a dedicated recording room. Upon completion, rats were deeply anesthetized with isoflurane (4%) and pentobarbital (3 ml, Pentoject, 200 mg/ml) and transcardially perfused with phosphate-buffered saline (PBS) followed by fixative (paraformaldehyde dissolved in PBS, 4%, wt/vol). Brains were extracted, sectioned and visualised to verify electrode location. Lesion severity was verified using antibodies to tyrosine hydroxylase (rabbit anti-TH primary; diluted 1:1000; Millipore Cat# AB152, RRID:AB_390204; donkey anti-rabbit Alexa Fluor 488 secondary; diluted 1:1000; Thermo Fisher Scientific Cat# A-21206, RRID:AB_2535792) to visualise the unilateral loss of cell bodies in the SNc and loss of dopaminergic innervation in the dorsal striatum. Only rats with stimulation electrode tips histologically verified as located in or adjacent to the border of globus pallidus were included in the reported data.

### Data acquisition and electrical stimulation

Electrocorticogram (ECoG) signals measured from M1.4 screws referenced to two cerebellar M1.4 screws were amplified and digitised using a RHD2000 family amplifying and digitising headstage connected to a RHD USB Interface Board both from Intan Technologies (intantech.com). Electrical stimulation was driven using a fully isolated current source (battery powered with an optically coupled trigger, A365RC, World Precision Instruments) with the source and sink terminals connected to the adjacent strands of the pair of stainless steel wires forming the stimulation electrode. Stimulation events were biphasic consisting of two consecutive pulses of opposite polarity each 50, 60 or 70 μA in amplitude and 95 μs in duration separated by 10 μs. To achieve closed-loop stimulation, a custom designed digital circuit was used to access the digital data stream from the headstage, track phase within a band of interest in real time and generate digital pulses to activate the optically coupled trigger of the stimulation current source. The custom designed digital circuit was implemented using extra available digital circuitry within the field programmable gate array (FPGA) located on the RHD USB Interface Board.

### Real-time phase tracking and closed-loop trigger algorithm

A novel approach was developed to track phase within a band of interest in real time. A continuous transform was performed directly on the digital values produced by the sampling analogue to digital converter (ADC). A sample rate of 20 kHz per channel was used in order to record the full detail of the stimulation artefacts aiding their removal in post hoc analysis. However, for phase tracking, the data was downsampled by producing a single sample from the sum of 8 to 12 (n_dn_) successive samples, since tracking beta-band phase did not require such a high number of samples per cycle. Thus, the phase tracking component ran at sample rates in the range 1.67 to 2.5 kHz (varying the sample rate was one of the ways to achieve different target oscillation centre frequencies). Performing a continuous transform allowed a phase estimate to be instantaneously produced corresponding to each value of the downsampled data stream. A bandlimited complex estimate of the signal was generated from the downsampled wideband data stream by, on receipt of each sample, updating a complex weight register that maintained the relationship between the estimate of the signal and a pair of continuously oscillating reference sine and cosine waves. The error between the real part of the estimate signal and the current ADC sample was used to update the weight register for use with the next ADC sample. The centre frequency of the band of interest was set by the frequency of the reference waves. This depended on the downsampled sample rate and the number of values in the lookup table containing a quarter of a sine wave used to generate the reference waves (n_points_, 11 to 15 values per quarter cycle were used). Thus the centre frequency of the phase tracker pass band was given by (f_c_ = 20000/n_dn_/n_points_/4). The width of the band of interest was set by a constant coefficient on the error term used to update the weight register. This essentially smoothed the reference wave by dampening the rate of convergence between the estimate and the signal. Finally, a continuous phase estimate was produced by calculating the polar angle of the complex signal estimate using a CORDIC (Volder, 1959). A real-time trigger was generated when the phase estimate passed into the target phase range unless it had been less than 0.8 of a target frequency period since the most recent passing into the target phase range. This was important to reduce stimulation at times of poor phase estimates caused by low power in the target frequency range. A phase shift corresponding to half the stimulation width (100 μs) was subtracted from the phase estimate causing the stimulation to be triggered early allowing the middle of the stimulation to occur at the desired target phase. Additionally, to reduce the effect of electrical stimulation artefacts on the phase estimate, a digital hold circuit operating on the 20 kHz data stream held the sample preceding the stimulation trigger on the input of the phase tracking circuit for 600 μs from the start of the trigger. An offset removal digital filter was also implemented on the input to the phase tracker to remove any residual offset remaining from the analogue front end.

### Recording protocol

Three to six recording blocks were performed per rat. Recording blocks consisted of eight pairs of recordings with a different real-time target phase applied in each pair. Target phase order was randomised across blocks. Stimulation was enabled in the first recording of each pair (stimulation recordings) and disabled in the second (baseline recordings). Stimulation recordings were approximately 4.5 minutes in duration and baseline recordings were approximately 2 minutes in duration. The real-time system generated triggers during 20 second epochs (on epochs) separated by 5 second trigger free epochs (off epochs) and was in operation across all recordings. Generating real-time triggers in the absence of closed-loop stimulation during baseline recordings allowed comparison of the real-time system performance between open-loop and closed-loop operation. These recordings also provided the opportunity to assess baseline physiology in the absence of stimulation. Embedding short epochs lacking stimulation within stimulation recordings provided an internal comparison point to assess the effect of stimulation without the need to correct for changes occurring over longer time frames such as arousal, brain state, general movement levels and external electrical noise. The centre frequency (f_c_) for real-time phase tracking for each block was chosen at the start of the block based on power spectra generated from previous recordings from the same rat and was in the range 34 to 38 Hz.

### Data analysis

Post hoc analysis of ECoG spectral properties and stimulation phase including statistical tests were performed using SciPy (scipy.org). Text values are reported as mean ± standard deviation and error bars in plots show mean ± standard error of the mean unless otherwise stated. First stimulation artefacts were removed from the 20 kHz recordings by interpolation between the sample immediately preceding the stimulation and the sample 1.7 ms later. The resulting signals were downsampled to 1 kHz in two steps using finite impulse response anti-aliasing filters (designed using remez, combined pass band ripple less than 0.001 dB below 420 Hz, stop band attenuation greater than 90 dB above 498 Hz). All spectra and phase calculations were performed on the artefact removed and downsampled signal. Post hoc trigger phase in the beta-band was calculated by filtering (firwin, numtaps = 513) the signal between ±5 Hz of the centre frequency (f_c_) chosen for real-time phase tracking. The trigger phase was then calculated as the polar angle of the Hilbert transform analytic signal at the midpoint of the biphasic trigger. Power spectral density (PSD) calculations were performed using Welch’s method (Welch, 1967) with a resolution of 1 Hz spectral bins. Changes in beta-band power were calculated frm the respective PSDs (mean of f_c_ bin ± 5 bins inclusive, i.e. mean of 11 bins total) in dB. The Watson-Williams test was performed using pycircstat (pypi.org/project/pycircstat).

